# The role of multilevel selection in host microbiome evolution

**DOI:** 10.1101/663351

**Authors:** Simon van Vliet, Michael Doebeli

## Abstract

Animals are associated with a microbiome that can affect their reproductive success. It is therefore important to understand how a host and its microbiome coevolve. According to the hologenome concept, hosts and their microbiome form an integrated evolutionary entity, a holobiont, on which selection can potentially act directly. However, this view is controversial and there is an active debate on whether the association between hosts and their microbiomes is strong enough to allow for selection at the holobiont level. Much of this debate is based on verbal arguments, but a quantitative framework is needed to investigate the conditions under which selection can act at the holobiont level. Here we use multilevel selection theory to develop such a framework. We found that selection at the holobiont level can in principle favor a trait that is costly to the microbes but that provides a benefit to the host. However, such scenarios require rather stringent conditions. The degree to which microbiome composition is heritable decays with time, and selection can only act at the holobiont level when this decay is slow enough, which occurs when vertical transmission is stronger than horizontal transmission. Moreover, the host generation time has to be short enough compared to the timescale of the evolutionary dynamics at the microbe level. Our framework thus allows us to quantitatively predict for what kind of systems selection could act at the holobiont level.

## Introduction

Most multicellular organisms are associated with a microbiome that can strongly affect their health (1). For example, microbes residing in the animal gut can help digest food, provide essential nutrients, condition the immune system, and can even affect the mental health of their host (2–5). As a result, the reproductive success of a host often depends on the composition of its microbiome (6). To understand the evolution of multicellular organisms it is thus important to understand how they coevolve with their microbiomes.

Because of these strong interdependencies some researchers have suggested that a host and its microbiome should be viewed as a single evolving entity, a so called holobiont (7–11). According to this hologenome concept, selection can potentially act directly at the level of the holobiont (7). As a result, traits that increase the overall reproductive success of a holobiont could evolve even if they are disfavored by selection at the level of the microbes.

For selection to act at the level of the holobiont it is essential that there is an association between the genotype of a host and the composition of its microbiome (12–14). Such an association could result from vertical transmission of the microbiome, i.e. when hosts pass a sample of their microbiome on to their offspring (11, 15, 16). This vertical transmission creates heritability in microbiome compositions allowing hosts with successful microbiomes to pass them on to their offspring. However, most hosts constantly take up microbes from their environment, weakening the heritable association between a host and its microbiome (11, 13, 15, 16). The pervasiveness of such horizontal transmission has led several researchers to conclude that selection at the holobiont level is unlikely to play a major role in nature (13, 14, 17–19).

The strength of the association between a host and its microbiome is thus expected to depend on the relative importance of vertical and horizontal transmission and recent studies have shown that organisms can vary strongly in this regard (11, 15). This raises the question for what organisms and under what conditions selection can act at the level of the holobiont. This question has been actively debated in the recent literature, however most of this debate is based on verbal arguments (7–10, 13, 17, 18). In a recent model the effects of horizontal and vertical transmission where studied for microbiomes consisting of only a single genotype (11, 20). However, by excluding variation in microbiome composition this model could not address the question under what conditions selection at the holobiont level can maintain traits that are disfavored by selection at the microbe level. This question is one of the most contentious points in the ongoing debate and we address it here using a mathematical model.

We investigated under what conditions microbes can evolve a trait that provides a benefit to the host but that comes at a cost to themselves. Such a trait is disfavored by selection at the microbe level and can thus only evolve when selection can act at the holobiont level. We used a multilevel selection framework to address this question, as this allowed us to independently control the strength and direction of selection at both the microbe and host level. This multilevel selection framework does not explicitly incorporate the holobiont concept, however asking under what conditions *holobiont level* selection can maintain a trait that is opposed by microbe level selection is equivalent to asking when *host level* selection can maintain such a trait. Using simulations we show that such a trait can evolve, but only when vertical transmission is stronger than horizontal transmission and when the host generation time is short enough compared to the timescale of the evolutionary dynamics at the microbe level.

## Results

### A simple model for host-microbe evolutionary dynamics

We extended a previously developed multilevel selection framework (21) to investigate under what conditions selection can act at the host level. Throughout, we assumed the simplest possible dynamics to reduces the number of model parameters. We consider a microbiome consisting of two genotypes: helper cells and neutral cells (Fig. 1). Helper and neutral cells are identical except for one trait: helper cells increase the reproductive success of their host, but this comes at a cost to their own growth rate. We assume that microbes have a constant birth rate of *β* for the neutral cells and (1 − *γ*)*β* for the helper cells, where *γ* is the cost of helping. We expect that within a given host, microbes can grow to a maximal population size and thus assume that death rates are density dependent (Fig. 1). Helper cells can transition into neutral cells and vice versa due to mutations. At each microbial division event there is thus a probability *µ* that a microbe mutates to the other type. Finally, we assume that microbes can only grow within the host environment and we thus only keep track of the microbial densities within each host. Microbiomes are characterized by the frequency of helper cells *f*_*i*_ (*helper frequency* in short). In particular we often show the mean helper frequency in the total host population 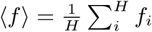.

**Fig. 1.**
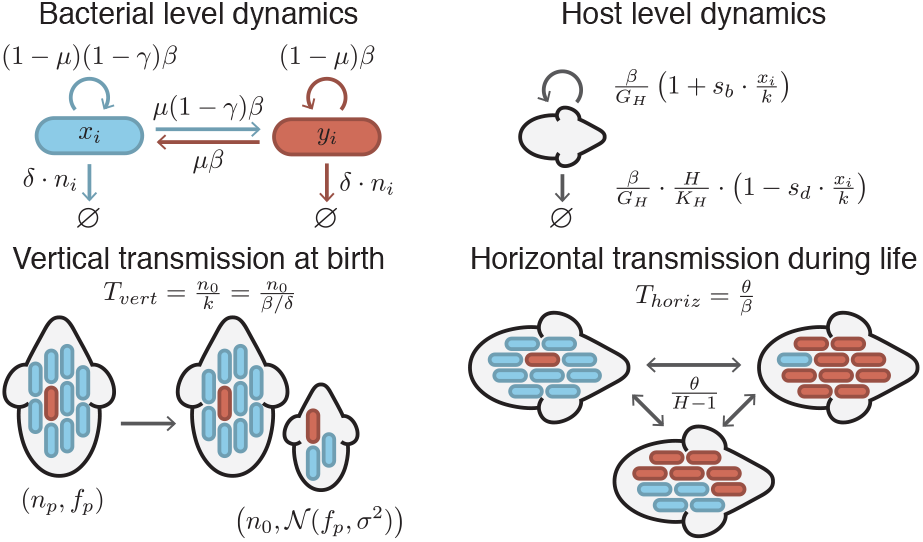
We consider a population of *H* hosts each carrying a microbiome consisting of two types: helper *x*_*i*_ and neutral *y*_*i*_ cells. These types are identical, except that helper cells pay a cost *γ* for increasing the host reproductive success. Newborn hosts are colonized by a sample of their parents microbiome, which has fixed density *n*_0_ and a frequency of helper cells drawn from a truncated normal distribution centered on the parent’s helper frequency *f*_*p*_ and with variance *σ*^2^. The strength of this vertical transmission 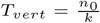 is given by the ratio of the size of this sample to the microbial carrying capacity 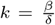. There is a constant exchange of microbes between hosts with rate *θ*. The strength of horizontal transmission 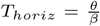 is given by the ratio of the migration rate to the microbial birth rate. Microbes have a constant birth rate of (1 − *γ*)*β* for helper and *β* for neutral cell, a density dependent death rate *δ* ⋅ *n*_*i*_, and a constant mutation rate *µ* between helper and neutral cells. Hosts have a birth rate that increase linearly with the density of helper cells (with slope *s*_*b*_) and a death rate that decreases linearly with density of helper cells (with slope *s*_*d*_), moreover the death rate increases linearly with the number of hosts. *G*_*H*_ measures the number of microbe generations per host generation. Microbiomes are described by their density *n*_*i*_ = *x*_*i*_ + *y*_*i*_ and frequency of helper cells *f*_*i*_ = *x*_*i*_/*n*_*i*_.

Helper cells increase the reproductive success of their host, for example by producing an essential nutrient or by providing protection against pathogenic bacteria. As a result helper cells can either increase host birth rates or decrease host death rates. We assume that host birth rates increase linearly with the density of helper cells from *β/G*_*H*_ in the absence of helper cells to (1 + *s*_*b*_) *β/G*_*H*_ when a host is fully occupied by helper cells. The parameter *s*_*b*_ thus controls how strong host birth rates depend on their microbiome. Similarly, we assume that host death rates decrease linearly with the density of helper cells, with the parameter *s*_*d*_ controlling the strength of this dependance (Fig. 1). Throughout the main text, we only consider the case where helper cells increase host birth rates without affecting host death rates (i.e. we assume *s*_*d*_ = 0). However, we obtained similar results when helper cells decreased host death rates instead (Fig SI2). To maintain bounded host population sizes we assume that hosts death rates are density dependent (Fig. 1). The generation time of hosts and microbes typically differ strongly, and as a result we expect evolutionary change to happen at different timescales for hosts and microbes. The parameter *G*_*H*_ in our model controls how many bacterial generations occur within each host generation.

We explicitly incorporate the two modes of transmission, vertical and horizontal, in our model (Fig. 1). When a host gives birth, it passes a sample of its microbiome on to its offspring. The strength of vertical transmission 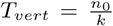 is measured as the size of this sample *n*0, relative to the microbial carrying capacity *k*. We expect that vertical transmission is subject to sampling variation: the helper frequency in a newborn hosts will likely differ slightly from that in their parent. We modeled sampling variation using a distribution in which we could independently control the degree of variance, allowing us to directly test the importance of sampling variation. Specifically, we assume that the helper frequency in a newborn host is drawn from a truncated normal distribution centered on the helper frequency of the parent and with a constant variance of *σ*^2^. Throughout their lives hosts exchange bacteria by horizontal transmission. We assume that bacteria migrate between hosts at a constant rate and that migration is random (i.e., no spatial structure). The strength of horizontal transmission 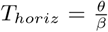 is measured as the migration rate *θ* relative to the bacterial growth rate *β*.

### Host level selection can maintain helper cells

Microbes do not receive any direct benefit from helping their host: helper cells grow slower than neutral cells but have the same death rate, consequently microbiomes consisting of only helper cells also reach a lower steady state density than microbiomes consisting of only neutral cells. Helper cells are thus expected to go extinct over evolutionary time in the absence of host level selection. Indeed our model shows that helper cells rapidly decrease in frequency when host birth rates are independent of microbiome composition (*s*_*b*_ = 0, Fig. 2B). Helper cells are only maintained at a low frequency due to constant transitions from neutral cells (i.e. at a frequency set by mutation-selection balance).

**Fig. 2.**
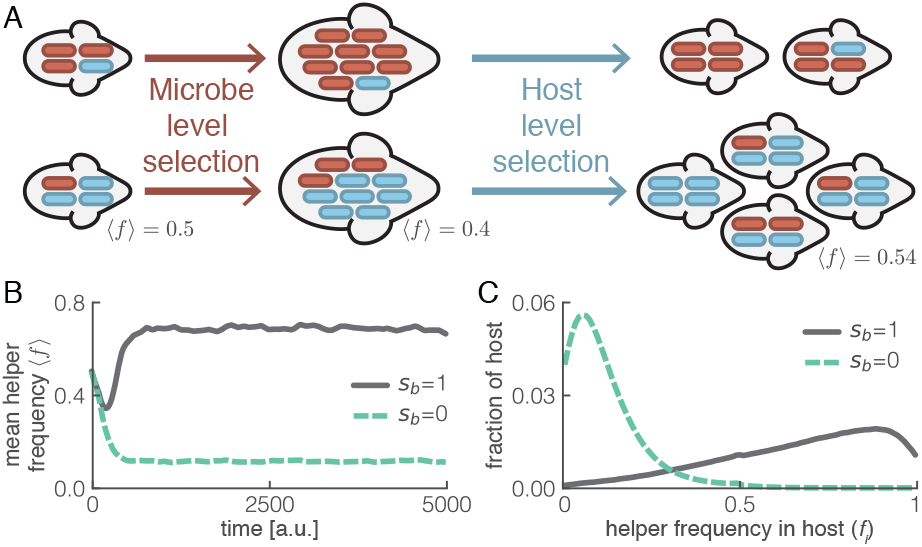
Selection at host level can maintain helper cells. (A) Within each host, helper cells (blue) decrease in frequency, because microbe level selection favors the faster growing neutral cells (red). However helper cells can increase in frequency due to host level selection: hosts with more helper cells have more offspring. These offspring in turn also have high helper frequencies because of vertical transmission. (B) Helper cells are only maintained at high frequencies when host birth rates depend on microbiome composition (*s*_*b*_ = 1); the average frequency of helper cells over the total host population (*mean helper frequency* ⟨*f* ⟩) is shown. (C) Frequency of helper cells *f*_*i*_ varies widely between hosts; the distribution is shown for t=5000. Parameters as shown in Table S1, except *G*_*H*_ = 10 and *K*_*H*_ = 5000.

Helper cells could potentially be maintained by host level selection when they increase host reproductive success (or decrease host death rates). The frequency of helper cells decreases within each host due to selection at the microbe level, however the frequency of helper cells within the total host population could increase because hosts with many helper cells have more offspring and pass on their microbiome (Fig. 2A). Indeed our model shows that helper cells can increase to high frequencies when host birth rates depend on the microbiome composition (*s*_*b*_ = 1, Fig. 2BC).

### Host level selection can only act under strong constraints

We performed an extensive exploration of the model parameter space to investigate when selection at the host level can maintain a trait that is disfavored by selection at the microbe level (Fig. SI1). We found that two conditions have to be met (Fig. 3): i) vertical transmission *T*_*vert*_ has to be strong compared to horizontal transmission *T*_*horiz*_; and ii) the host generation time *τ*_*H*_ has to be short enough compared to the timescale *τ*_*M*_ of the evolutionary dynamics at the microbe level (i.e. the timescale over which helper frequencies decrease within a single host). These two conditions can be understood as follows: when vertical transmission is weak compared to horizontal transmission all hosts will quickly converge to same microbiome composition. As a result there is no longer any variation in helper frequencies between hosts and without this variation there can be no host level selection. Likewise, if helper cells within a host have gone extinct before a host can reproduce (due to microbe level selection), there is again no variation between hosts for selection to act on.

**Fig. 3.**
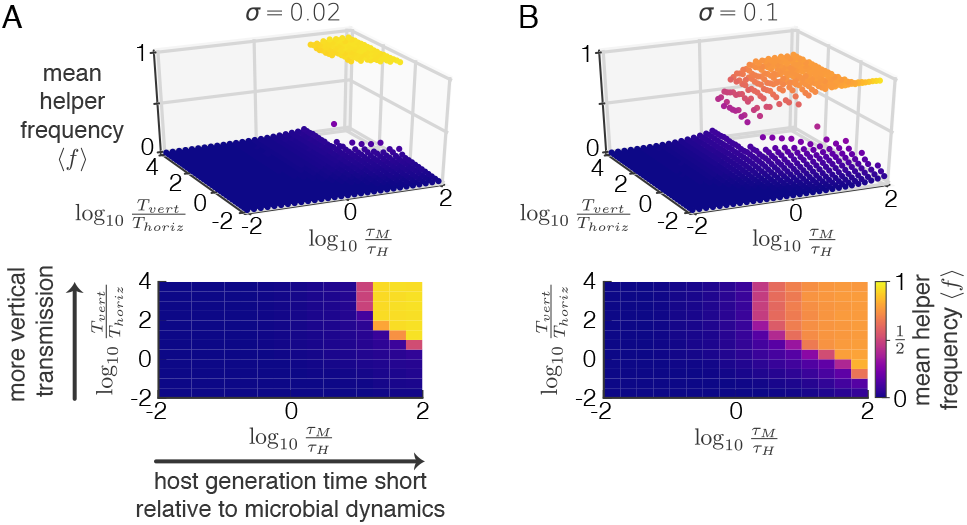
Selection can only act at the host level under stringent conditions. The mean helper frequency is shown as function of the timescale of the evolutionary dynamics at the microbe level 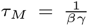 relative to the host generation time 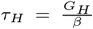 and the ratio of vertical 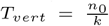 to horizontal transmission 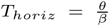. Two different levels of sampling variance are shown, the standard deviation of the sampling distribution *σ* is 0.02 in (A) and 0.1 in (B); top panels show the result of single simulations, bottom panels show the same data, but averaged within discrete bins. We varied the cost *γ* and migration rate *θ* to obtain different values of *τ*_*M*_/*τ*_*H*_ and *T*_*vert*_/*T*_*horiz*_, respectively, all other parameters as shown in Table S1.

Two additional factors are important in determining when host level selection can maintain helper cells: the region of parameter space in which helper cells can reach high frequencies is larger when the sampling variance is higher (Fig. 3) and when the host birth rates depends more strongly on the microbiome composition (Fig. SI1). In the next two sections we will explore these requirements in more detail.

### Heritability of microbiome composition decays with time

For host level selection to maintain helper cells, the microbiome composition needs to be hereditable. At birth, a host is colonized by vertically transmitted cells, but during its life it acquires cells by horizontal transmission at a constant rate (Fig. 4A). Consequently, the proportion of vertically transmitted cells decreases over time (Fig. 4B). The microbiome of a host thus shifts from being dominated by microbes obtained from its parent to being dominated by microbes obtained from the environment (i.e. the other hosts in the population). In other words, the degree to which community composition is heritable decreases over time.

**Fig. 4.**
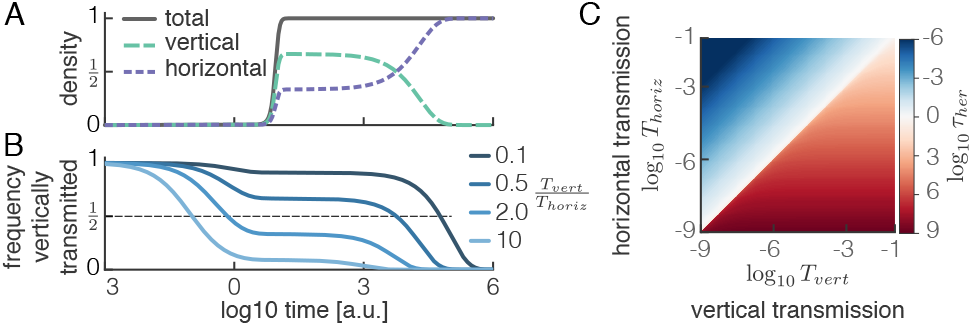
Microbiome composition is only heritable when vertical transmission dominates horizontal transmission. (A) Horizontally acquired cells eventually dominate over vertically acquired cells. The density of the total population and of vertically and horizontally acquired cells was calculated analytically (SI eq. 7,8) and is shown for the case *T*_*vert*_ = 2*T*_*horiz*_. (B) Frequency of vertically transmitted cells decreases over time in a non-linear way. The frequency of vertically inherited cells was calculated analytically (SI eq. 8) and is shown for four different ratios of vertical to horizontal transmission. Microbiome composition is heritable (i.e. dominated by vertically transmitted cells) for long periods when vertical transmission dominates (dark blue) and for short periods when horizontal transmission dominates (light blue). (C) Heritability timescale *τ*_*her*_ is short when vertical transmission *T*_*vert*_ is weak compared to horizontal transmission *T*_*horiz*_. The heritability timescale *τ*_*her*_ is defined as the time after birth at which the frequency of vertically acquired cells reaches 0.5 and was calculated numerically (SI eq. 8).

We define the heritability timescale *τ*_*her*_ as the time after host birth at which the frequency of cells that entered the host by vertical transmission reaches 50%. In other words, when *t* > *τ*_*her*_ the microbiome composition has become dominated by microbes obtained from the environment. We used our model to calculate this timescale numerically and we derived an analytical approximation (SI Appendix 1). We found that it depends in a non-linear way on the relative strength of horizontal and vertical transmission (Fig. 4BC). This nonlinearity is caused by the non-linear growth dynamics of the microbes: horizontally acquired cells that arrive directly after birth change the microbiome composition to a much larger extent than those arriving when the microbiome has reached its steady state density. When horizontal transmission is dominant (*T*_*horiz*_ > *T*_*vert*_) the microbiome composition resembles that of the environment long before the microbiome reaches its steady state density and heritability is rapidly lost (*τ*_*her*_ < 1, Fig. 4BC). In contrast, when vertical transmission dominates, the microbiome reaches its steady state density before horizontally acquired cells can take over. As a result, heritability can be maintained over long time periods that are proportional to the inverse of the migration rate (*τ*_*her*_ ∝ 1/*θ*, Fig. 4BC, SI Appendix 1).

### Maintenance of helper cells

Within a single host, helper cells always decrease in frequency due to microbe level selection. We can analytically show that this decrease follows a sigmoidal curve with a timescale that is inversely proportional to the cost of helping: *τ*_*M*_ = 1/(*βγ*) (Fig 5A, SI Appendix 2). For helper cells to be maintained by host level selection, hosts have to give birth before all helper cells are lost. The host generation time *τ*_*H*_ = *G*_*H*_/*β* thus has to be short enough compared to the timescale *τ*_*M*_ of the evolutionary dynamics at the microbe level (Fig 5B). The bigger the difference in generation time between hosts and microbes, the lower the cost of helping has to be in order to maintain helper cells (Fig 5B).

**Fig. 5.**
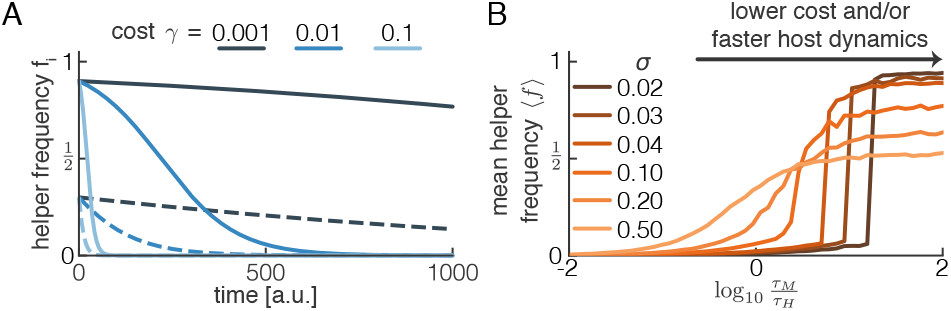
Host generation time has to be short enough compared to the timescale of the evolutionary dynamics at the microbe level. (A) Within a given host the frequency of helper cells decreases following a sigmoidal curve with timescale *τ*_*M*_ = 1/(*γβ*). The frequency of helper cells was calculated analytically (SI eq. 22) for an isolated host (no migration) and is shown for three different costs *γ* starting from an initial frequency of 0.9 (solid) or 0.3 (dashed). (B) Host level selection can only maintain helper cells when the host generation time *τ*_*H*_ = *G*_*H*_/*β* is short enough compared to the timescale of the evolutionary dynamics at the microbe level *τ*_*M*_. This ratio *τ*_*M*_/*τ*_*H*_ = 1/(*γG*_*H*_) depends on the cost of cooperation *γ* and number of microbe generations per host generation *G*_*H*_. Higher sampling variance *σ*^2^ allows for host level selection over a larger region of parameter space, but lowers the maximal value of the helper frequency. We varied the cost *γ* to obtain different values of *τ*_*M*_/*τ*_*H*_, all other parameters as shown in Table S1.

Sampling variation is an essential component in our model, as it is the only process that can increase the frequency of helper cells over evolutionary time. Within a host the helper frequency always decreases; it is lower at the time when a host reproduces than when it was born. Without sampling variation each subsequent generation of hosts would thus inherit a helper frequency that is lower than the helper frequency of its parents at the time of their birth. Over many host generations helper cells would thus disappear. Sampling variation can counteract this process because it can create some offspring that have helper frequencies that are higher than that of their parents. To maintain helper cells over evolutionary time, sampling variation has to be strong enough to offset the decrease in helper frequency that occurred during a parent’s lifetime in a sufficiently large number of offspring. The higher the cost of helping, the more the helper frequency decreases during a parent’s lifetime, and the higher the sampling variation has to be in order to maintain helper cells (Fig 5B). When sampling variation is high and the cost of helping is low, helpers can be maintained at intermediate frequencies even when microbiome composition is not heritable (i.e. when horizontal transmission is stronger than vertical transmission, Fig 3B). However, in this case helper cells are not maintained by host level selection, but as result of sampling variance: helper cells achieve the same frequency when host birth rates are independent of microbiome composition (Fig. SI3).

High sampling variation allows host level selection to maintain helper cells in a larger region of parameter space, but it lowers the maximally achieved helper frequency (Fig 5B). This is because the helper frequency is bounded between 0 and 1. When a parent already has a high helper frequency, the frequency in its offspring can increase by only a small amount but can decrease by a large amount. As a result the distribution of helper frequencies in newborn hosts becomes skewed towards lower helper frequencies and this decreases its average value. In regions of parameter space where the average helper frequency is high, increasing sampling variance increases the skew in the distribution of helper frequencies in newborn hosts. As a result, the average helper frequency in the total population has a lower steady state value in this region (Fig 3B, SI1). For a similar reason increasing sampling variance will increase the average helper frequency in regions of parameter space where the average helper frequency is low (Fig 3B, SI1). Although sampling variation at host birth is essential in our model, other mechanisms can in principle be envisaged to maintain helper cells. The essential requirement for such mechanisms is that they can increase the frequency of helper cells in newborn hosts compared to their parents. For example, higher rates of vertical transmission of helper cells compared to neutral cells could also generate such a scenario.

### Helping behavior can evolve *De novo*

So far we have assessed under what conditions host level selection can maintain pre-existing microbial types that provide a costly benefit to their host. However, we have not yet addressed the question of how such costly traits could evolve in the first place. Here we develop a second model to address this question.

We consider a single microbial species characterized by a continuous trait, which is called the *cooperative investment*, that determines how much help is provided to the host. The birth rate of the microbes decreases linearly with the level of investment (i.e. the cost increases linearly), while the birth rate of the hosts increases linearly with the total cooperative investment of its microbiome. Every time a microbe divides there is small probability that its offspring will mutate to a slightly higher or lower investment level. Otherwise, the model dynamics are the same as in the two cell-type model discussed previously (see methods for details).

The conditions under which high cooperative investment levels can evolve are similar to the ones required to maintain helper cells: vertical transmission needs to be strong compared to horizontal transmission and the cost of cooperation has to be low compared to the inverse of the host generation time (*γ <* 1/*G*_*H*_, Fig SI4). When these conditions are met, cooperative investment levels can increase rapidly to close to their maximal value of 1 (Fig. 6A). The distribution of investment levels within the global microbial population is sharply peaked with most microbes having an investment level between 0.8 and 1 (Fig. 6B). As a result of host level selection, microbes can thus evolve to express a trait that benefits their host, but that reduces their own growth rate.

**Fig. 6.**
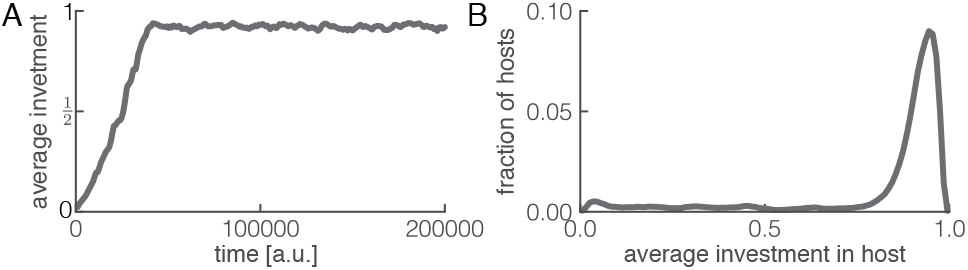
Costly microbial traits can evolve de novo by host level selection. Microbes have a continuous trait, called the *cooperative investment*; microbial growth rates decrease linear with investment while host growth rates increase linearly with the total cooperative investment of its microbiome. (A) Average level of cooperative investment in the global microbial population increases rapidly over time. (B) The distribution of cooperative investment levels at steady state is sharply peaked at high investment levels. Parameters as shown in table S2.

## Discussion

We found that traits that are costly to a microbe, but beneficial to their host, can evolve by selection at the host level, but only under stringent conditions. We derived these conditions using simple dynamics for both hosts and microbes, however the framework we developed can easily be extended to incorporate more complex dynamics at either level. Moreover, we expect that our conclusions are qualitatively robust to relaxing many of our assumptions. Although we considered only a two species microbiome, our results would also hold in a more complex system, as long as the other members of the microbiome interact in an equal way with helper and neutral cells. Moreover, selection at the level of the host always requires that microbes providing the benefit have not gone extinct by the time that the host replicates and we can show (SI Appendix 2) that this timescale generally depends on the cost of cooperation. Finally, the degree to which the community composition is heritable generally decays with time as long as there is non-zero horizontal transmission (SI Appendix 1).

Throughout, we assumed that hosts are passive players. Many hosts, however, do strongly interact with their microbiome (19): the immune system can control the growth of microbes (22) and hosts can reward helpful microbes (23). Such interactions could facilitate host level selection. For example, horizontal transmission could be much higher when hosts can filter incoming migrants based on their identity (24). Likewise, hosts could offset the cost incurred by the microbes by providing directed benefits to helper cells (23). Our framework can readily be extended to incorporate such host-microbe interactions, providing interesting opportunities for future work. In our current model we ignored host effects to get insights into when host level selection can act by itself. By extension, this also reveals when host effects are likely to be essential. Moreover, we expect that in many biological systems, hosts could not tell the difference between helper and neutral cells, for example because they only differ by a small number of mutations. In this scenario the host would interact identically with both helper and neutral cells and our model would directly apply, as in this case all host effects could be absorbed by adjusting our model parameters.

Our model suggests that selection at the host level, and in extension at the holobiont level, could be of importance in nature, but only under rather restrictive circumstances: host generation times have to be short enough relative to the timescale of the evolutionary dynamics at the microbe level and vertical transmission has to dominant over horizontal transmission. These conditions could potentially be met in certain short lived insect species, but are unlikely to be fulfilled in long lived mammals. For most species, host level effects would be essential to allow for selection at the level of the holobiont. Although other authors have come to similar conclusions (13, 14, 17–19), the lack of a quantitative framework prevented direct predictions for when host level selection could be of importance. The mathematical framework we developed here provides some tools to start exploring how host and microbiomes coevolve in natural settings.

## Methods

### Microbial dynamics

We consider a population of *H*(*t*) hosts and model the density of helper cells *x*_*i*_(*t*) and neutral cells *y*_*i*_(*t*) in each host *i* using differential equations. The per capita birth rate is *β* for neutral cells and (1 − *γ*)*β* for helper cells, where *γ* is the cost of providing a benefit to the host. At each microbe birth event there is a small probability *µ* that helper cells mutate into neutral cells and vice versa. The per capita death rate is *δ ⋅ n*_*i*_(*t*) for both helper and neutral cells, where *n*_*i*_(*t*) = *x*_*i*_(*t*) + *y*_*i*_(*t*) is the total microbial density in host *i*. With these assumptions, the microbial dynamics are given by:

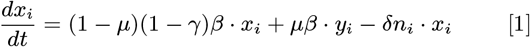

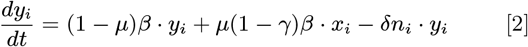

### Host dynamics

Hosts dynamics are modeled using an individual based approach. The birth rate *B*_*i*_ of host *i* increases linearly with the density of helper cells from a basal level of 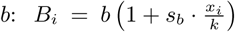, the normalization of helper cell densities by the microbial carrying capacity *k* = *β/δ* ensures that host birth rates remain bounded between *b* and *b* + *s*_*b*_. The death rate rate *D*_*i*_ increases linearly with the number of hosts and decreases linearly with the density of helper cells in the host: 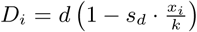 The constants *s*_*b*_ and *s*_*d*_ control how strongly host reproductive success depends on microbiome composition, and thus control the strength of selection at the host level. Throughout the main text we use *s*_*d*_ = 0. It is convenient to rewrite these rates as:

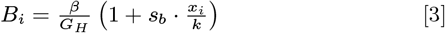

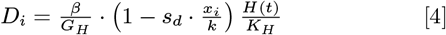

where *G*_*H*_ = *β/b* is the number of microbial generations per host generation and *K*_*H*_ = *b/d* is the expected number of hosts (host carrying capacity) at steady state when *s*_*b*_ = *s*_*d*_ = 0.

### Vertical transmission

When a new host is born, it is seeded with a microbiome of fixed total density *n*_0_ and a helper frequency 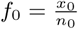 which is drawn from a truncated normal distribution with constant variance *σ*^2^ and a mean value equal to the helper frequency in its parent *f*_*p*_:

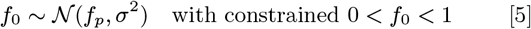

### Horizontal transmission

Microbes leave a host at a fixed rate *θ* and are distributed evenly among all other hosts in the population. The inflow of helper cells into host *i* is thus given by: 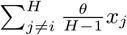, where *θ* ⋅ *x*_*j*_ is the number of helper cells leaving hosts *j* and 1/(*H* − 1) is the fraction of those cells ending up in host *i*. As horizontal transmission acts continuously, we modify the dynamical equations for the microbiome dynamics (eq. 1 and 2) by including the terms for the in and outflow of microbes:

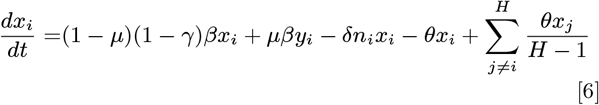

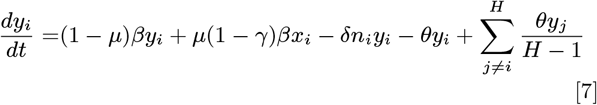

### Model parameters

We can reduce the number of independent parameters in our model by measuring time in units of the inverse microbial birth rate 1/*β* and by measuring microbial densities in units of their carrying capacity *k* = *β/δ*. In all simulations we therefore arbitrarily set *β* = *δ* = *k* = 1. See Table S1 for all other parameter values used in the simulations.

### Model implementation

The model was solved numerically using code implemented in Python. We updated the microbial densities and host birth and death rates using a coarse time step ∆*t* and used a fine time step *δt* ≤ ∆*t* to implement host birth and death events assuming constant rates. We used this procedure to keep computation times reasonably short and confirmed that we obtain identical results if we update microbial dynamics and hosts rates every *δt* time step. At each time step ∆*t* we performed the following steps:

1. Calculate birth (*B*_*i*_) and death (*D*_*i*_) rates for each host.
2. Iterate with time step *δt*; in each *δt* time step at maximum a single host level event (birth or death) can occur. If there are *H* hosts in the populations, there are 2*H* + 1 possible events that can happen in this time step: any of the *H* hosts can reproduce, host *i* reproduces with probability *P* (*b*_*i*_) = *B*_*i*_ ⋅ *δt*; any of the *H* hosts can die, host *i* dies with probability *P* (*d*_*i*_) = *D*_*i*_ ⋅ *δt*; or no host event happens, with probability 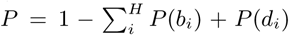. We randomly select one of these possible events based on their relative probabilities. If a host gives birth we add a new host to the population with a microbiome of density *n*_0_ and helper frequency *f*_0_ given by eq. 5. Newborn hosts cannot give birth or die until the next ∆*t* time step. If a host dies we remove that host and its microbiome from the population.
3. Update microbiome state using Euler method: *x*_*t*+1_ = *x*_*t*_ + ∆*t ⋅ g*_*x*_(*x*_*t*_, *y*_*t*_) and *y*_*t+1*_ = *y*_*t*_ + ∆*t* ⋅ *g*_*y*_ (*x*_*t*_, *y*_*t*_), where *g*_*x*_ and *g*_*y*_ are given by the right hand site of eq. 6 and 7 respectively.

The time step *δt* was dynamically adjusted to the number of hosts in the population such that the probability of having more than one host event per time step was less than 0.01. We ran the simulations until the helper frequency reached its steady state level; this condition was evaluated automatically by requiring that the variation in helper frequency over a moving time window (10^4^ time units) was below a threshold value. All simulation results were averaged over a moving time window of 1000 time units.

### Continuous investment model

Next we consider a model where the degree to which a microbe helps its host is a continuous trait, called the cooperative investment 0 < *w* < 1. To solve the model we discretized the investment level into *N* = 100 equally sized bins; in each bin *j* the investment is given by *w*^*j*^ = (2*j* − 1)/(2*N*), *j* = (1, 2,…, *N*). We track 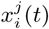, the density of bacteria with investment level *j* in host *i*, over time. Each host can thus be characterized by the total microbial density 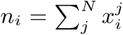 and investment distribution 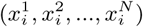. Bacteria with investement level *w*^*j*^ have a birth rate of (1 − *w*^*j*^ ⋅ *γ*)*β*. With probability (1 − *µ*) their offspring inherits the same investment level, and with probability *µ*/2 their offspring mutates to an investment level of *w*^*j*^ ± *ϵ*. We set *ϵ* = 1/*N* so that mutations only happen between adjacent investment bins. There is a density dependent death rate *δ* ⋅ *n*_*i*_ and constant migration between hosts with rate *θ*. The dynamics for 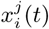 are then given by:

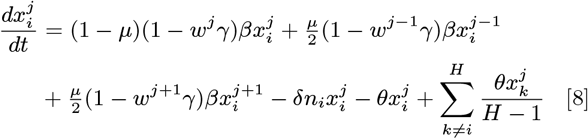

where 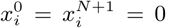. The host birth and death rate are given by:

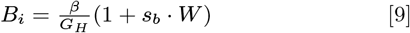

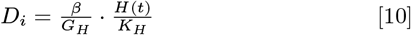

where 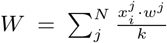 is the cumulative investment of all microbes in host *i* (the normalization by the microbial carrying capacity *k* insures that 0 *< W <* 1). Vertical transmission is implemented by randomly drawing *N*_0_ discrete samples from the parent investment distribution with probabilities 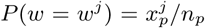. The resulting offspring investment distribution is normalized to a total density of *n*_0_. The model is implemented identically to the two-type model described above, but now using eq. 8 to update microbiome state and eqs. 9 and 10 to calculate host rates.

## Supporting information

Supplemental Infromation

## Author contributions

SvV: Conceptualization, Methodology, Formal analysis, Writing - original draft. MD: Conceptualization, Supervision, Writing - review & editing.

## Acknowledgements

We thank Alma Dal Co, Gil Henriques and the other members of the Doebeli lab for comments and discussion. SVV was supported by SNF Postdoc.Mobility fellowship 175123 and MD was supported by NSERC Discovery grant 219930.

## References

1. McFall-Ngai M, et al. (2013) Animals in a bacterial world, a new imperative for the life sciences. Proc. Natl. Acad. Sci. U. S. A. 110(9):3229–36.

2. Honda K, Littman DR (2016) The microbiota in adaptive immune homeostasis and disease. Nature 535(7610):75–84.

3. Schroeder BO, Bäckhed F (2016) Signals from the gut microbiota to distant organs in physiology and disease. Nat. Med. 22(10):1079–1089.

4. Postler TS, Ghosh S (2017) Understanding the Holobiont: How Microbial Metabolites Affect Human Health and Shape the Immune System. Cell Metab. 26(1):110–130.

5. Vuong HE, Yano JM, Fung TC, Hsiao EY (2017) The Microbiome and Host Behavior. Annu. Rev. Neurosci. 40(1):21–49.

6. Gould AL, et al. (2018) Microbiome interactions shape host fitness. Proc. Natl. Acad. Sci. 115(51):E11951–E11960.

7. Zilber-Rosenberg I, Rosenberg E (2008) Role of microorganisms in the evolution of animals and plants: the hologenome theory of evolution. FEMS Microbiol. Rev. 32(5):723–735.

8. Bordenstein SR, Theis KR (2015) Host Biology in Light of the Microbiome: Ten Principles of Holobionts and Hologenomes. PLOS Biol. 13(8):e1002226.

9. Theis KR, et al. (2016) Getting the Hologenome Concept Right: an Eco-Evolutionary Framework for Hosts and Their Microbiomes. mSystems 1(2):e00028–16.

10. Rosenberg E, Zilber-Rosenberg I (2018) The hologenome concept of evolution after as Units of Selection and a Model of Their Population Dynamics and Evolution. Biol. Theory 13(1):44–65.

11. Roughgarden J, Gilbert SF, Rosenberg E, Zilber-Rosenberg I, Lloyd EA (2018) Holobionts as Units of Selection and a Model of Their Population Dynamics and Evolution. Biol. Theory 13(1):44–65.

12. Godfrey-Smith P (2009) Darwinian Populations and Natural Selection, Darwinian Populations and Natural Selection. (OUP Oxford).

13. Douglas AE, Werren JH (2016) Holes in the Hologenome: Why Host-Microbe Symbioses Are Not Holobionts. MBio 7(2):e02099–15.

14. Stencel A, Wloch-Salamon DM (2018) Some theoretical insights into the hologenome theory of evolution and the role of microbes in speciation. Theory Biosci. 137(2):197–206.

15. Engel P, Moran NA (2013) The gut microbiota of insects diversity in structure and function. FEMS Microbiol. Rev. 37(5):699–735.

16. Moeller AH, Suzuki TA, Phifer-Rixey M, Nachman MW (2018) Transmission modes of the mammalian gut microbiota. Science (80-.). 362(6413):453–457.

17. Moran NA, Sloan DB (2015) The Hologenome Concept: Helpful or Hollow?. PLOS Biol. 13(12):e1002311.

18. Skillings D (2016) Holobionts and the ecology of organisms: Multi-species communities or integrated individuals? Biol. Philos. 31(6):875–892.

19. Foster KR, Schluter J, Coyte KZ, Rakoff-Nahoum S (2017) The evolution of the host microbiome as an ecosystem on a leash. Nature 548(7665):43–51.

20. Roughgarden J (2018) Holobiont Evolution: Model for Vertical vs. Horizontal Microbial Colonization. bioRxiv p. 465310.

21. Simon B, Fletcher JA, Doebeli M (2013) Towards a general theory of group selection. Evolution (N. Y). 67(6):1561–1572.

22. Brown EM, Sadarangani M, Finlay BB (2013) The role of the immune system in governing host-microbe interactions in the intestine. Nat. Immunol. 14(7):660–667.

23. Schluter J, Foster KR (2012) The Evolution of Mutualism in Gut Microbiota Via Host Epithelial Selection. PLoS Biol. 10(11):e1001424.

24. Kaltenpoth M, et al. (2014) Partner choice and fidelity stabilize coevolution in a Cretaceous-age defensive symbiosis. Proc. Natl. Acad. Sci. 111(17):6359–6364.

