## Supplemental Infromation for "The role of multilevel selection in host microbiome evolution"

#### This PDF file includes:

Supplementary text
Figs. S1 to S4
Tables S1 to S2

### Supporting Information Text

#### Appendix 1 — Timescale of loss of heritability of microbiome composition

When horizontal transmission rates are non-zero, over time the microbiome of a given host transitions from a state where it is dominated by cells acquired by vertical transmission to a state dominated by cells acquired by horizontally transmission. As a result the heritability of microbiome composition decays over time. Here we develop a simple model to quantify the timescale over which heritability is lost. We focus on a single genotype and track the density of cells acquired by vertical transmission $x^v$ and of cells acquired by horizontal transmission $x^h$. The dynamics of these two groups are identical, except for the mode of immigration. We can write down the following general equations for the dynamics of vertically and horizontally acquired cells in host $i$:

$$\frac{dx_i^v}{dt} = \left[b(n_i) - d(n_i) - m^{out}(n_i)\right] \cdot x_i^v, \qquad x_i^v(0) = n_0 \tag{1}$$

$$\frac{dx_i^h}{dt} = \left[b(n_i) - d(n_i) - m^{out}(n_i)\right] \cdot x_i^h + m_i^{in}(t), \qquad x_i^h(0) = 0 \tag{2}$$

where $b(n)$, $d(n)$, and $m_{out}(n)$ are arbitrary functions describing the per capita birth, death, and emigration rates, and $m_i^{in}(t)$ is the (potentially time dependent) rate of immigration into host $i$. As vertically and horizontally acquired cells are identical except for their origin, these rates only depend on the total density $n_i = x_i^v + x_i^h$.

**General solution.** We are primarily interested in the frequency of vertically transmitted cells $f_i^v = \frac{x_i^v}{x_i^v + x_i^h}$ and can rewrite equations 1 and 2 as:

$$\frac{df_i^v}{dt} = -\frac{m_i^{in}}{n_i(t)} \cdot f_i^v, \qquad f_i^v(0) = 1 \tag{3}$$

$$\frac{dn_i}{dt} = \left[b(n_i) - d(n_i) - m^{out}(n_i)\right] \cdot n_i + m_i^{in}(t), \qquad n_i(0) = n_0 \tag{4}$$

We can integrate equation 3 to find an implicit solution for $f_i^v(t)$:

$$f_i^v(t) = e^{-\int_0^t \frac{m_i^{in}(\tau)}{n_i(\tau)} d\tau} \tag{5}$$

The frequency of vertically transmitted cells thus decays exponentially with a time dependent rate that depends on the rate of immigration relative to the current population size.

**Specific solution.** Equation 5 holds for all possible rate functions, however to find an explicit solution we need to specify the rate functions. We follow the assumptions stated in the main text and assume constant birth and migration rates and a density dependent death rate:

$$b(n_i) = \beta, \qquad d(n_i) = \delta \cdot n_i, \qquad m^{out}(n_i) = \theta, \qquad m_i^{in}(t) = \tfrac{\theta}{H-1} \sum_{i \neq j} n_j(t)$$

As host dynamics are typically much slower than microbiome dynamics, we further assume that microbial densities in all other hosts in the population are at their steady state level: $n_j(t) = k \; \forall i \neq j$, where $k = \frac{\beta}{\delta}$ is microbial carrying capacity. With this assumption the rate of immigration simplifies to:

$$m_i^{in}(t) = \theta \cdot k$$

With these assumptions eq. 4 becomes:

$$\frac{dn}{dt} = \left[\beta - \theta - \beta \cdot \tfrac{n}{k}\right] \cdot n + \theta \cdot k, \qquad n(0) = n_0 \tag{6}$$

We have dropped the subscript $i$ as we only consider the dynamics in the focal host. The analytical solution for $n(t)$ is given by:

$$n(t) = k \cdot \frac{\left(n_0 + k\frac{\theta}{\beta}\right) \cdot e^{(\theta+\beta)t} - \frac{\theta}{\beta}(k - n_0)}{\left(n_0 + k\frac{\theta}{\beta}\right) \cdot e^{(\theta+\beta)t} + (k - n_0)} \tag{7}$$

From eq. 5 we can then find the frequency of vertically inherited cells:

$$f^v(t) = \frac{n_0(1+\frac{\theta}{\beta})\cdot e^{\beta t}}{(n_0+k\frac{\theta}{\beta})\cdot e^{(\theta+\beta)t} - \frac{\theta}{\beta}(k-n_0)} \quad [8]$$

We now define the heritability timescale $\tau_{her}$ as the time when the frequency of vertically transmitted cells reaches 0.5: $f^v(\tau_{her}) = \frac{1}{2}$. Although we have an explicit expression for $f^v(t)$, we cannot analytically solve for the heritability time. However we can find the numerical solution (Fig. 4C), and in the next section we will derive an analytical approximation.

**Analytical approximation.** The non-linear density dependent death rate in eq. 6 makes it impossible to solve eq. 8 analytically for the heritability timescale. Here we combine two approximations to derive an analytical expression for the heritability timescale. The first approximation applies to the early phase of host colonization, where microbial densities are low ($\frac{n}{k} \ll 1$); the second approximation applies to the late phase of colonization, where the microbial density is close to the steady state value ($n \approx k$). For both regimes we derive an expression for $f^v(t)$. We then combine these approximations at the time where microbial densities reach half their carrying capacity ($n = \frac{k}{2}$) to obtain an approximation for the entire system.

In the early phase we assume that $\frac{n}{k} \ll 1$ and ignore the density dependent death rate in eq. 6:

$$\frac{dn}{dt} \approx (\beta-\theta)\cdot n + \theta \cdot k \qquad n(0) = n_0. \quad [9]$$

We can solve this equation for $n_{early}(t)$, the approximate microbial density early during colonization:

$$n_{early}(t) = \frac{(n_0(\beta-\theta)+k\theta)\cdot e^{(\beta-\theta)t} - k\theta}{\beta-\theta}. \quad [10]$$

Using eq. 5 we can then find the approximate frequency of vertically transmitted cells during colonization:

$$f^v_{early}(t) = \frac{1}{1-\frac{k\theta}{n_0(\beta-\theta)}\cdot(1-e^{-(\beta-\theta)t})} \quad [11]$$

For the late phase of colonization we assume that $n(t) \approx k$, with this assumption equation 3 becomes:

$$\frac{df^v_{late}}{dt} = -\theta \cdot f^v_{late}. \quad [12]$$

The initial condition of the differential equation can be derived by requiring continuity between the approximations for the early and late phases of the colonization. We set the transition point at $n = \frac{k}{2}$. From eq. 10 we calculate the time $t_{1/2}$ when the microbial density reaches half its carrying capacity $n_{early}(t = t_{1/2}) = \frac{1}{2}$:

$$t_{1/2} = \frac{1}{\beta-\theta}\log\left[\frac{k(\beta+\theta)}{2\,(n_0(\beta-\theta)+k\theta)}\right] \quad [13]$$

From eq. 11 we then find the frequency of vertically transmitted cells at that time:

$$f_{1/2} \equiv f^v_{early}(t = t_{1/2}) = \frac{n_0(\beta+\theta)}{n_0(\beta-\theta)+k\theta} \quad [14]$$

We thus solve eq. 12 with the initial condition $f^v_{late}(t_{1/2}) = f_{1/2}$:

$$f^v_{late}(t) = \frac{n_0}{k}\cdot 2^{\frac{-\theta}{\beta-\theta}}\cdot\left(\frac{k(\beta+\theta)}{n_0(\beta-\theta)+k\theta}\right)^{\frac{\beta}{\beta-\theta}}\cdot e^{-\theta t} \quad [15]$$

Combining the two approximation, equations 11 and 15, we thus find:

$$f^v(t) \approx \begin{cases} \frac{1}{1-\frac{k\theta}{n_0(\beta-\theta)}\cdot(1-e^{-(\beta-\theta)t})} & \text{if } t \leq t_{1/2} \\ \frac{n_0}{k}\cdot 2^{\frac{-\theta}{\beta-\theta}}\cdot\left(\frac{k(\beta+\theta)}{n_0(\beta-\theta)+k\theta}\right)^{\frac{\beta}{\beta-\theta}}\cdot e^{-\theta t} & \text{if } t > t_{1/2} \end{cases} \quad [16]$$

We now calculate the approximate heritability timescale by solving $f^v(\tau_{her}) = \frac{1}{2}$ for $\tau_{her}$:

$$\tau_{her} \approx \begin{cases} \frac{1}{\beta-\theta}\log\left[\frac{k\theta}{\theta(k+n_0)-n_0\beta}\right] & \text{if } \frac{\theta}{\beta} > \frac{n_0}{k-3n_0} \\ \frac{1}{\beta-\theta}\log\left[\frac{k(\beta+\theta)}{2(n_0(\beta-\theta)+k\theta)}\right] + \frac{1}{\theta}\log\left[\frac{2n_0(\beta+\theta)}{n_0(\beta-\theta)+k\theta}\right] & \text{if } \frac{\theta}{\beta} \leq \frac{n_0}{k-3n_0} \end{cases} \quad [17]$$

Typically $n_0 \ll k$ and $\theta \ll \beta$, so we can simplify these equations further to:

$$\tau_{her} \approx \begin{cases} \frac{1}{\beta}\log\left[\frac{\theta/\beta}{\theta/\beta-n_0/k}\right] & \text{if } \frac{\theta}{\beta} > \frac{n_0}{k-3n_0} \\ \frac{1}{\theta}\log\left[\frac{2n_0/k}{n_0/k+\theta/\beta}\right] & \text{if } \frac{\theta}{\beta} \leq \frac{n_0}{k-3n_0} \end{cases} \quad [18]$$

These equation can be rewritten using the definitions for the strength of vertical transmission $T_{vert} = \frac{n_0}{k}$ and horizontal transmission $T_{horiz} = \frac{\theta}{\beta}$ to:

$$\tau_{her} \approx \begin{cases} \frac{1}{\beta}\log\left[\frac{T_{horiz}}{T_{horiz}-T_{vert}}\right] & \text{if } T_{horiz} > \frac{T_{vert}}{1-3T_{vert}} \\ \frac{1}{\theta}\log\left[\frac{2T_{vert}}{T_{vert}+T_{horiz}}\right] & \text{if } T_{horiz} \leq \frac{T_{vert}}{1-3T_{vert}} \end{cases} \quad [19]$$

We thus see that when horizontal transmission is stronger than vertical transmission ($T_{horiz} > T_{vert}$), heritability is quickly lost. Specifically, when $T_{horiz} > \frac{e}{e-1} \cdot T_{vert}$, heritability is lost on a timescale faster than the timescale over which the microbes grow ($\tau_H < \frac{1}{\beta}$). On the other hand, when horizontal transmission is much weaker than vertical transmission ($T_{horiz} \ll T_{vert}$), then heritability can be maintained over long timescales set by the migration rate: $\tau_H \approx \frac{\log 2}{\theta}$.

### Appendix 2 — Timescale of the evolutionary dynamics at the microbe level

Helper cells pay a cost for helping their host and will thus be replaced by the faster growing neutral cells in a single host. Here we will derive an expression for the timescale over which microbe level selection will cause helper cells to decrease in frequency.

We consider a single isolated host and ignore host level birth and death events. We assume that there are no transitions between helper and neutral cells (i.e. $\mu = 0$) and that helper and neutral cells are identical, except that helper cells pay a constant cost for helping their host that reduces their own birth rate. Under these assumptions we can write the following differential equations for the density of helper $x(t)$ and neutral cells $y(t)$:

$$\frac{dx}{dt} = (1-\gamma)\cdot b(x,y)\cdot x - d(x,y)\cdot x \quad [20]$$

$$\frac{dy}{dt} = b(x,y)\cdot y - d(x,y)\cdot y \quad [21]$$

where $\gamma$ is the fixed cost of helping and $b(x,y)$ and $d(x,y)$ are functions that describe the potentially state dependent per capita birth and death rates. From eq. 20 and 21 it follows that the frequency of helper cells $f = \frac{x}{x+y}$ changes over time as:

$$\frac{df}{dt} = -\gamma \cdot b(f,n)\cdot f(1-f) \quad [22]$$

where $n = x + y$ is the total population size. To solve for the frequency of helper cells we need to specify the birth rate function. Following the main text, we assume that birth rates are constant: $b(x,y) = b(f,n) = \beta$; the solution of eq. 22 is then given by:

$$f(t) = \frac{f_0 \cdot e^{-\gamma\beta\cdot t}}{f_0\,(e^{-\gamma\beta\cdot t}-1)+1} \quad [23]$$

where $f_0 \equiv f(t=0)$ is the initial frequency of helper cells. The frequency of helper cells thus decreases over time following a sigmoidal curve with timescale $\tau_M = \frac{1}{\gamma\beta}$.

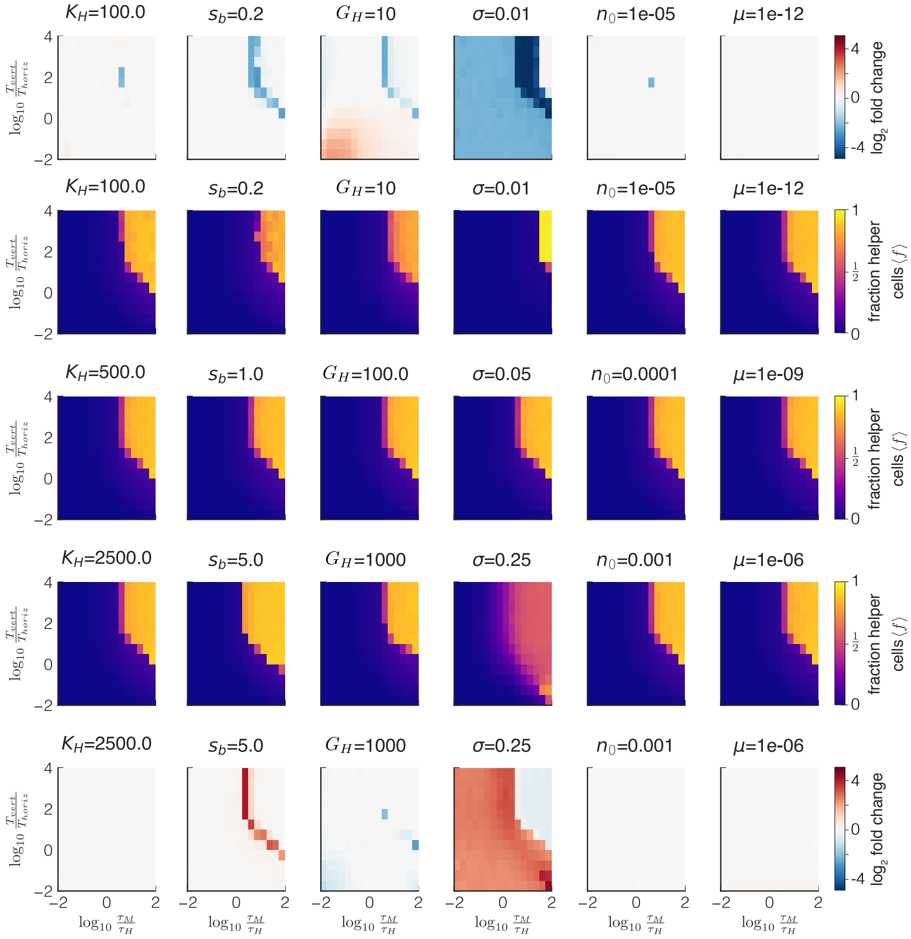


**Fig. S1.** Parameter sensitivity of the model. The center row shows the average helper frequency $\langle f \rangle$ as a function of the timescale of the evolutionary dynamics at the microbe level $\tau_M$ relative to the host generation time $\tau_H$ (obtained by varying the cost $\gamma$) and the ratio of vertical $T_{vert}$ to horizontal $T_{horiz}$ transmission (obtained by varying the migration rate $\theta$) for the default parameters; the row above and below the middle show the average helper frequency when the indicated parameter is changed; the top and bottom rows show the fold difference (log2 transformed) between the changed and default parameter value. Increasing the strength of selection $s_b$ increases the parameter regime in which host level selection can maintain helper cells; increasing the sampling variance $\sigma$ also increases the parameter space in which helper cells can be maintained, but it comes at the cost of lowering the maximal obtained helper frequency. All other parameters do not greatly affect the parameter space in which helper cells can be maintained. All other parameter values as shown in Table S1.

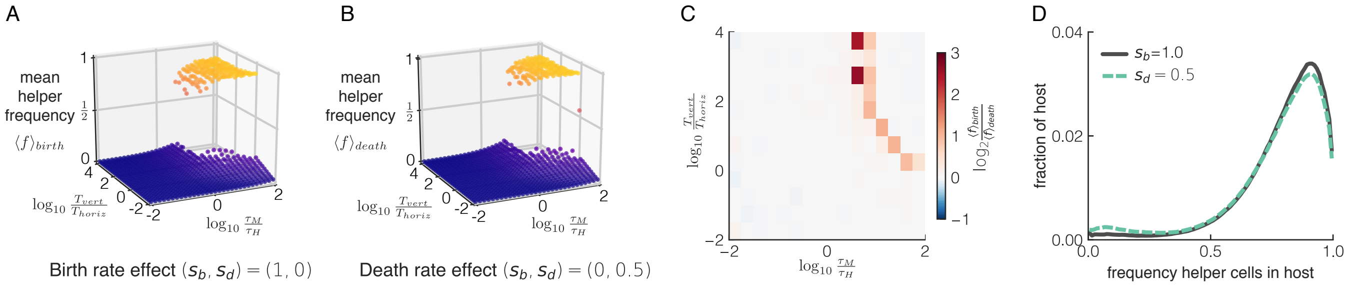


**Fig. S2.** Similar dynamics are obtained when helper cells affect birth or death rates. We compare two models: in model I helper cells increase host birth rates, but do not effect host death rates $s_b = 1$, $s_d = 0$; in model II helper cells decrease host death rates, but do not effect host birth rates $s_b = 0$, $s_d = 0.5$; the values of $s_b$ and $s_d$ are chosen such that the expected number of hosts is the same for the two models when $\langle f \rangle = 0$ and when $\langle f \rangle = 1$. (A,B) Average helper frequency $\langle f \rangle$ as a function of the timescale of the evolutionary dynamics at the microbe level $\tau_M$ relative to the host generation time $\tau_H$ (obtained by varying the cost $\gamma$) and the ratio of vertical $T_{vert}$ to horizontal $T_{horiz}$ transmission (obtained by varying the migration rate $\theta$) for model I (A) and model II (B). (C) Fold difference (log2 transformed) in helper frequency between model I and model II. (D) Distribution of helper frequency within the host population for model I (solid grey) and model II (dashed green), for the case where $\tau_M/\tau_H = 10$ and $T_{vert}/T_{horiz} = 100$. When helper cells increase host birth rates (model I) host level selection can maintain them in a slightly larger parameter regime than when helper cells lower death rates (model II), but the overall dynamics are very similar. Parameter values as shown in Table S1.

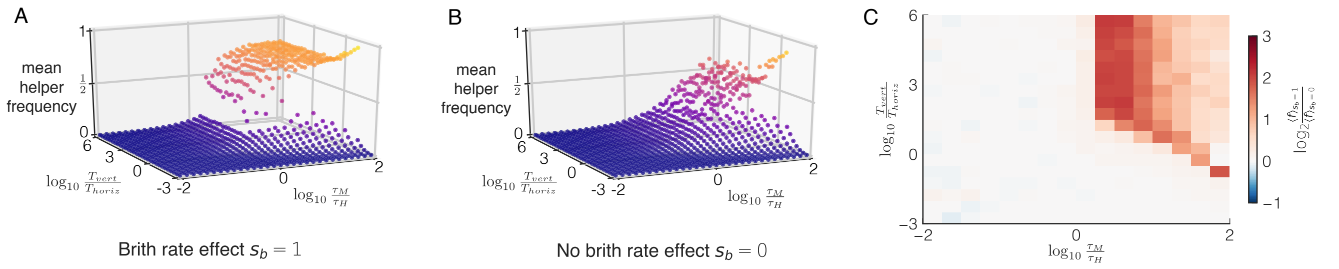


**Fig. S3.** Sampling variation can maintain helper cells at intermediate frequencies in absence of selection. We compare two models: in model I helper cells increase host birth rates $s_b = 1$. In model II host fitness is independent of microbiome composition, $s_b = 0$, and thus the same for all hosts. As a result, there is no selection at the level of the host in model II. (A,B) Average helper frequency $\langle f \rangle$ as a function of the timescale of the evolutionary dynamics at the microbe level $\tau_M$ relative to the host generation time $\tau_H$ (obtained by varying the cost $\gamma$) and the ratio of vertical $T_{vert}$ to horizontal $T_{horiz}$ transmission (obtained by varying the migration rate $\theta$) for model I (A) and model II (B). (C) Fold difference (log2 transformed) in helper frequency between model I and model II. When the cost of helping is low ($\tau_M > \tau_H$) and sampling variation is high ($\sigma = 0.1$) helper cells can be maintained by sampling variation even in the absence of host level selection. Parameter values as shown in Table S1, except for $\sigma = 0.1$.

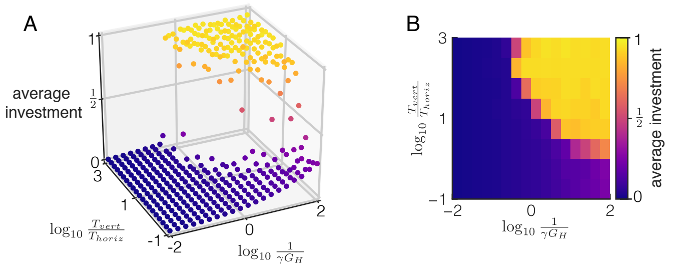


**Fig. S4.** De novo evolution of costly helping behavior requires vertical transmission to dominate over horizontal transmission and the number of microbial generations per host generation $G_H$ to be small compared to the inverse cost of helping $\gamma$. Microbes have a continuous trait value, the cooperative investment, that determines how much they help their host. Birth rates of microbes decrease linearly with the investment level of the microbe and birth rates of the host increase linearly with the total investment of the microbiome. (A,B) The average evolved level of investment is shown as function of $\frac{1}{\gamma G_H}$ and the ratio of vertical $T_{vert}$ to horizontal $T_{horiz}$ transmission. $\frac{1}{\gamma G_H}$ is equivalent to $\frac{\tau_M}{\tau_H}$ in the two species model and measures the timescale of the evolutionary dynamics at the microbe level ($\propto \beta/\gamma$) relative to the host generation time ($\propto G_H/\beta$). (A) Shows results of individual simulations (B) shows the same data, but averaged within discrete bins. We varied the cost $\gamma$ and migration rate $\theta$ to obtain different values of $1/(\gamma G_H)$ and $T_{vert}/T_{horiz}$, respectively, all other parameters as shown in Table S2.

**Table S1. Parameters of two-type (helper and neutral cell) model. The simulations were run using the default value unless otherwise stated in the figure captions. To vary $\tau_M/\tau_H = 1/(\gamma G_H)$ we changed the value of $\gamma$ while keeping $G_H$ constant. To vary $T_{vert}/T_{horiz} = (\beta n_0)/(k\theta)$ we changed $\theta$ while keeping $n_0$ constant ($\beta$ and $k$ can be arbitrarily set to 1 and were thus never varied). We obtained identical results when we varied both $\gamma$ and $G_H$ to obtain different values of $\tau_M/\tau_H$ and when we varied both $\theta$ and $n_0$ to obtain different values of $T_{vert}/T_{horiz}$. We verified that our results were robust to the time step $\Delta t$: we obtained similar results when we used smaller time steps.**

| **Parameter Name** | **Symbol** | **Default Value** | **Notes** |
|---|---|---|---|
| Microbe birth rate | $\beta$ | 1 | Arbitrarily set to 1 (measure time in units of $1/\beta$) |
| Microbe death rate | $\delta$ | 1 | Arbitrarily set to 1 (measure microbial densities in units of $k$) |
| Microbe mutation rate | $\mu$ | $10^{-9}$ | Mutation rate between helper and neutral cells |
| Cost of helping | $\gamma$ | 0.01 | |
| Microbe migration rate | $\theta$ | $10^{-6}$ | |
| Density of vertically transmitted sample | $n_0$ | $10^{-4}$ | |
| Standard deviation of sampling distribution | $\sigma$ | 0.05 | Sampling variance is $\sigma^2$ |
| Dependance of host birth rate on microbiome | $s_b$ | 1 | |
| Dependance of host death rate on microbiome | $s_d$ | 0 | |
| Nr. of microbial generations per host generation | $G_H$ | 100 | |
| Host carrying capacity | $K_H$ | 500 | |
| Time step | $\Delta t$ | 0.05 | |
| | | **Derived quantities** | |
| **Parameter Name** | **Symbol** | **Relation** | |
| Microbial carrying capacity | $k$ | $\frac{\beta}{\delta}$ | |
| Strength of vertical transmission | $T_{vert}$ | $\frac{n_0}{k}$ | |
| Strength of horizontal transmission | $T_{horiz}$ | $\frac{\theta}{\beta}$ | |
| Time scale of evolutionary dynamics at microbe level | $\tau_M$ | $\frac{1}{\beta\gamma}$ | |
| Host generation time | $\tau_H$ | $\frac{G_H}{\beta}$ | |

**Table S2. Parameters of the continuous investment model. The simulations were run using the default value unless otherwise stated in the figure captions. To vary $1/(\gamma G_H)$ we changed the value of $\gamma$ while keeping $G_H$ constant. To vary $T_{vert}/T_{horiz} = (\beta n_0)/(k\theta)$ we changed $\theta$ while keeping $n_0$, $\beta$, and $k$ constant. We verified that our results were robust to the time step $\Delta t$: we obtained similar results when we used smaller time steps.**

| Parameter Name | Symbol | Default Value | Notes |
|---|---|---|---|
| Microbe birth rate | $\beta$ | $1$ | Arbitrarily set to 1 (measure time in units of $1/\beta$) |
| Microbe death rate | $\delta$ | $1$ | Arbitrarily set to 1 (measure microbial densities in units of $k$) |
| Microbe mutation rate | $\mu$ | $0.01$ | Mutation rate between different cooperative investment levels |
| Maximum cost of helping | $\gamma$ | $0.01$ | |
| Microbe migration rate | $\theta$ | $10^{-5}$ | |
| Density of vertically transmitted sample | $n_0$ | $10^{-3}$ | |
| Number of investment levels sampled | $N_0$ | $10$ | |
| Dependance of host birth rate on microbiome | $s_b$ | $1$ | |
| Nr. of microbial generations per host generation | $G_H$ | $100$ | |
| Host carrying capacity | $K_H$ | $500$ | |
| Time step | $\Delta t$ | $0.05$ | |
| | | **Derived quantities** | |
| **Parameter Name** | **Symbol** | **Relation** | |
| Microbial carrying capacity | $k$ | $\frac{\beta}{\delta}$ | |
| Strength of vertical transmission | $T_{vert}$ | $\frac{n_0}{k}$ | |
| Strength of horizontal transmission | $T_{horiz}$ | $\frac{\theta}{\beta}$ | |
| Host generation time | $\tau_H$ | $\frac{G_H}{\beta}$ | |
